# Dynamic reconfiguration, fragmentation and integration of whole-brain modular structure across depths of unconsciousness

**DOI:** 10.1101/783175

**Authors:** Dominic Standage, Corson N. Areshenkoff, Joseph Y. Nashed, R. Matthew Hutchison, Melina Hutchison, Dietmar Heinke, Ravi S. Menon, Stefan Everling, Jason P. Gallivan

**Author notes:** Correspondence should be addressed to: Dominic Standage, School of Psychology, University of Birmingham, Jason Gallivan, Centre for Neuroscience Studies, Queen’s University. The authors declare no competing financial interests.

## Abstract

General anesthetics are routinely used to induce unconsciousness, and much is known about their effects on receptor function and single neuron activity. Much less is known about how these local effects are manifest at the whole-brain level, nor how they influence network dynamics, especially past the point of induced unconsciousness. Using resting-state functional magnetic resonance imaging (fMRI) with nonhuman primates, we investigated the dose-dependent effects of anesthesia on whole-brain temporal modular structure, following loss of consciousness. We found that higher isoflurane dose was associated with an increase in both the number and isolation of whole-brain modules, as well as an increase in the uncoordinated movement of brain regions between those modules. Conversely, we found that higher dose was associated with a decrease in the cohesive movement of brain regions between modules, as well as a decrease in the proportion of modules in which brain regions participated. Moreover, higher dose was associated with a decrease in the overall integrity of networks derived from the temporal modules, with the exception of a single, sensory-motor network. Together, these findings suggest that anaesthesia-induced unconsciousness results from the hierarchical fragmentation of dynamic whole-brain network structure, leading to the discoordination of temporal interactions between cortical modules.

## INTRODUCTION

The biological basis of consciousness has long fascinated neuroscientists, psychologists and clinicians alike. This fascination not only stems from the broad implications of consciousness on our understanding of human awareness and conscious experience, but also because disruptions in consciousness are a hallmark of both pathological (neurology) and naturally occurring physiological states (sleep), and are often purposefully induced in clinical treatment (anesthesiology). In this regard, the neuroscientific investigation of consciousness has been greatly advanced in recent decades by the study of the unconscious state. Considerable work now focuses on changes in brain states that accompany normal and pathological conditions like sleep and coma respectively (Mashour and Hudetz 2018). Likewise, the pharmacological induction and reversible manipulation of consciousness by general anesthesia has proven to be a productive line of investigation. Understandably, much of this latter work has focused on the cellular effects of anesthetics (Anis et al. 1983; Peduto et al. 1991; Wu et al. 1996), but it is unclear how these local effects are manifest as unconsciousness at the large-scale, global level (Franks 2006; Alkire et al. 2008; Brown et al. 2011).

According to integrated information theory (Tononi 2004), consciousness is linked to the ability to integrate information both locally and between widely distributed brain areas. Information integration is also a fundamental principle of global neuronal workspace theory [see (Dehaene et al. 2014)], which posits critical roles for long-distance interactions between brain regions in prefrontal, parietal and cingulate cortex during general awareness. These theories suggest that general anesthetics impair functional interactions across remote brain areas, causing the fragmentation of whole-brain networks (Mashour and Hudetz 2018). Integrated information theory further posits that consciousness is not a fixed state per se, but is graded in nature, decreasing proportionately with the number of discriminable brain states (Tononi et al. 2016). This gradient further suggests that interactions between brain networks (and consequently information integration) diminish beyond the initial threshold of unconsciousness (light sedation), with greater fragmentation at deeper levels of sedation. Thus, investigation of the dose-dependent effects of anesthesia on whole-brain networks not only quantifies the effects of sedation on the global properties of neural processing, but also serves to characterise the very nature of unconsciousness.

In recent years, resting-state functional MRI (rs-fMRI) has provided tremendous insight into the large-scale, network-level effects of anesthesia. The spontaneous, low-frequency signals recorded with rs-fMRI reflect underlying changes in neural activity and have been shown to correspond to known patterns of whole-brain connectivity in multiple species (Fox and Raichle 2007; Vincent et al. 2007; Shmuel and Leopold 2008; Hutchison and Everling 2012; Hutchison et al. 2012, 2015; Leopold and Maier 2012). Initially, the effects of anesthesia on network connectivity were largely focused on activity within selected networks and were assessed by constructing static functional networks from full scans (Vincent et al. 2007; Boveroux et al. 2010; Deshpande et al. 2010; Liu et al. 2011; Liu, Zhu, et al. 2013). Over these longer timescales, resting-state functional networks may appear similar during levels of wakefulness and anesthesia (Barttfeld et al. 2015), since static measurements of functional connectivity (FC) may largely recapitulate the underlying anatomical map (Honey et al. 2009; Hermundstad et al. 2013). There is a growing appreciation, however, that functional networks are temporally dynamic (Chang and Glover 2010; Bassett et al. 2011; Hutchison, Womelsdorf, Allen, et al. 2013; Hutchison, Womelsdorf, Gati, et al. 2013; Allen et al. 2014) and that an understanding of whole-brain computation requires time-resolved methods for network analysis (Medaglia et al. 2015; Bassett and Sporns 2017).

In the study of unconsciousness, changes in the repertoire of whole-brain states derived from functional networks constructed over short timescales (e.g., 30 s - 1.5 minutes) have been shown to correlate with anaesthetic dose (Hutchison et al. 2014; Barttfeld et al. 2015; Hudetz et al. 2015; Uhrig et al. 2018). Specifically, it has been reported that the number of discriminable brain states decreases at deeper levels of unconsciousness, consistent with predictions by integrated information theory (Tononi et al. 2016). Together with static FC analyses demonstrating the fragmentation of networks during unconsciousness (Achard et al. 2012)(Boly et al. 2012; Spoormaker et al. 2012; Monti et al. 2013; Hutchison et al. 2014), this finding implies that the temporal dynamics of network modularity may be highly informative. Modularity refers to the degree to which networks can be decomposed into groups of nodes (brain regions), with dense connectivity within these groups and sparse connectivity between them (formally quantifying fragmentation) (Sporns and Betzel 2016). Modular structure has long been recognised as a crucial property of complex biological systems, conferring functional specialisation and robustness to change (including damage) (Kirschner and Gerhart 1998; Kashtan and Alon 2005; Wagner 2013). Analyses of task-based changes to modular structure have been highly productive in recent studies of the whole-brain bases of motor skill acquisition (Bassett et al. 2011, 2015) and cognitive performance (Braun et al. 2015), but to the best of our knowledge, no studies have leveraged these methods in the study of unconsciousness (see the Discussion for a summary of earlier studies using static measures of modularity).

We examined the dose-dependent effects of isoflurane on the temporal modular structure of whole-brain networks in nonhuman primates, following induction of unconsciousness. We tested three principal hypotheses. First, we sought to determine whether whole-brain networks would show evidence of greater fragmentation at deeper levels of unconsciousness under our time-resolved approach. Thus, we tested the hypothesis that the number of temporal modules, and their degree of modularity, will increase with isoflurane dose. Second, we reasoned that if the strength of connectivity within brain networks decreases with increasing dose [e.g. (Boveroux et al. 2010; Guldenmund et al. 2013)], then smaller perturbations (e.g. background noise) may be sufficient to drive small, random network reconfigurations, leading to uncoordinated changes in modular membership. We therefore quantified the degree to which brain regions move between modules in a coordinated (and uncoordinated) manner at different levels of dose. Third, we sought to determine the extent to which dynamic whole-brain structure would break down in a spatially uniform vs. selective manner at deeper levels of unconsciousness. Thus, we derived networks from the temporal modules and tested the null hypothesis that the strength of interactions within and between those networks would decrease in a uniform fashion across networks with increasing isoflurane dose.

## METHODS

### Animal Preparation

Five macaque monkeys (Macaca fascicularis; four females; weights ranging from 3.6 to 5.3 kg [mean ± standard deviation = 4.26 ± 0.76 kg] and ages ranging from 7.71 to 8.22 years [mean ± standard deviation = 7.83 ± 0.22 years]), were involved in the experiment. All surgical and experimental procedures were carried out in accordance with the Canadian Council of Animal Care policy on the use of laboratory animals and approved by the Animal Use Subcommittee of the University of Western Ontario Council on Animal Care. Note that the use of an animal model for investigating the effects of anaesthesia on brain activity offers greater standardization between subjects and circumvents concerns of potentially inducing a lethal collapse of the cardiovascular or respiratory system at high dosage in humans. The complete methods for this experiment have previously been described in detail (Hutchison et al. 2014). We therefore provide a more concise description of the methods relevant to our temporal analyses.

Prior to image acquisition, monkeys were injected intramuscularly with atropine (0.4 mg/kg), ipratropium (0.025 mg/kg), and ketamine hydrochloride (7.5 mg/kg), followed by intravenous administration of 3 ml propofol (10 mg/ml) via the saphenous vein. Animals were then intubated and switched to 1.5% isoflurane mixed with medical air. Each monkey was then placed in a custom-built monkey chair and inserted into the magnet bore, at which time the isoflurane level was lowered to 1.00%. Prior to image localization, shimming, and echo-planar imaging (EPI), at least 30 min was allowed for the isoflurane level and global hemodynamics to stabilize at this 1.00% concentration. We then acquired 2 functional EPI scans at each of six increasing isoflurane levels: 1.00, 1.25, 1.50, 1.75, 2.00, and 2.75% (0.78, 0.98, 1.17, 1.37, 1.56, and 2.15 minimum alveolar concentration [MAC], respectively). We interleaved a 10 min period between each increase in isoflurane dose to allow for the concentration to stabilize. Throughout the duration of scanning, the monkeys spontaneously ventilated and we monitored physiological parameters (temperature, oxygen saturation, heart rate, respiration, and end-tidal CO_2_) to ensure that values were within normal limits. [See Supplementary Figure 1 in Hutchison et al., 2014]. The acquisitions of two anatomical images occurred during the stabilization periods between isoflurane dose levels.

### Data Acquisition

The monkeys were scanned on an actively shielded 7-Tesla 68-cm horizontal bore scanner with a DirectDrive console (Agilent, Santa Clara, California) with a Siemens AC84 gradient subsystem (Erlangen, Germany). We used a custom in-house conformal five-channel transceive primate-head Radio Frequency (RF) coil. Each functional run consisted of 150 continuous EPI functional volumes (repetition time [TR] = 2000 ms; echo time [TE] = 16 ms; flip angle = 70^0^; slices = 36; matrix = 96 x 96; Field of view [FOV] = 96 x 96 mm^2^; acquisition voxel size = 1 x 1 x 1 mm^3^), acquired with GRAPPA = 2. A high-resolution gradient-echo T2 anatomical image was acquired along the same orientation as the functional images (TR = 1100 ms, TE = 8 ms, matrix = 256 x 256, FOV = 96 x 96 mm^2^, acquisition voxel size = 375 x 375 x 1,000 mm^3^). We also acquired a T1-weighted anatomical image (TE = 2.5 ms, TR = 2300 ms, FOV = 96 x 96 mm^2^, acquisition voxel size = 750 x 750 x 750 mm^3^).

### Image Preprocessing and Analysis

Functional image preprocessing was implemented in the FMRIB Software Library toolbox (FSL; http://www.fmrib.ox.ac.uk). This consisted of motion correction (six-parameter affine transformation), brain extraction, spatial smoothing (Gaussian kernel of full-width at half maximum 3 mm applied to each volume separately), high-pass temporal filtering (Gaussian-weighted least-squares straight line fitting with sigma = 100 s), and low-pass temporal filtering (half-width at half maximum = 2.8 s, Gaussian filter). Functional data were nonlinearly registered to the T2 anatomical (FNIRT; http://www.fmrib.ox.ac.uk/fsl/fslwiki/FNIRT), and then registered to the T1 anatomical (six degrees of freedom rigid transformation), and finally normalized (12 degrees of freedom linear affine transformation) to the F99 atlas template (Van Essen 2004) [see http://sumsdb.wustl.edu/sums/macaquemore.do].

The F99-template normalized Lewis and van Essen (Lewis and Van Essen 2000a, 2000b) divisions were used to define 174 (87 per hemisphere) cortical regions (see Supplementary Figure S1 for an overview of these regions). Following regression of the average white matter (WM), cerebrospinal fluid (CSF) and six motion parameters from the regional time series, we calculated the mean time series for each region by averaging the time series across all voxels contained within it. Note that we did not use regions from subcortical structures, due to concerns about anatomically based parcellation of small substructures and decreased signal-to-noise ratio.

There is mounting evidence that variation in functional connectivity (FC) between regions and networks that occur during the resting state reflect real neural processes that are ignored in standard resting-state investigations that examine FC over the entire resting-state time window (Chang and Glover 2010; Hutchison, Womelsdorf, Allen, et al. 2013; Hutchison, Womelsdorf, Gati, et al. 2013; Allen et al. 2014). To explore the potential effects of isoflurane dose on time-varying network dynamics, each of the 174 regional time series was divided into windows of 60 seconds (30 imaging volumes) where contiguous windows overlapped by 50%. We constructed functional networks in each time window by calculating the Pearson correlation coefficient between each pair of brain regions (Figure 1C). All correlations that did not pass false discovery rate correction (Benjamini and Yekutieli 2001) were set to zero (q=0.05), as were all negative correlations (a requirement of the module detection approach used). We determined the modular structure of the resulting multilayer (a.k.a. multislice) networks with a generalized Louvain method for time-resolved clustering (Jeub et al, http://netwiki.amath.unc.edu/GenLouvain, 2011-2017). This algorithm was repeated 100 times with random initialisation, resulting in 100 clustering solutions (a.k.a. partitions). On each repetition, the algorithm was iterated until a stable partition was found, i.e. the output on iteration *n* served as the input on iteration *n*+1 until the output matched the input (Mucha et al. 2010; Bassett et al. 2011). In the main text, we used the standard spatial and temporal resolution parameters *γ* = 1 and ω = 1 respectively (see the Supplementary Material for demonstrations of robustness). Note that our choice of window length served a balance between temporal resolution and mitigation against the effects of noise on network construction (Sakoğlu et al. 2010; Hutchison, Womelsdorf, Allen, et al. 2013; Leonardi and Van De Ville 2015), while the overlap between windows served to increase the number of layers in the multi-layer networks. Neither parameter was crucial to our results (see the Supplementary Material).

**Figure 1:**
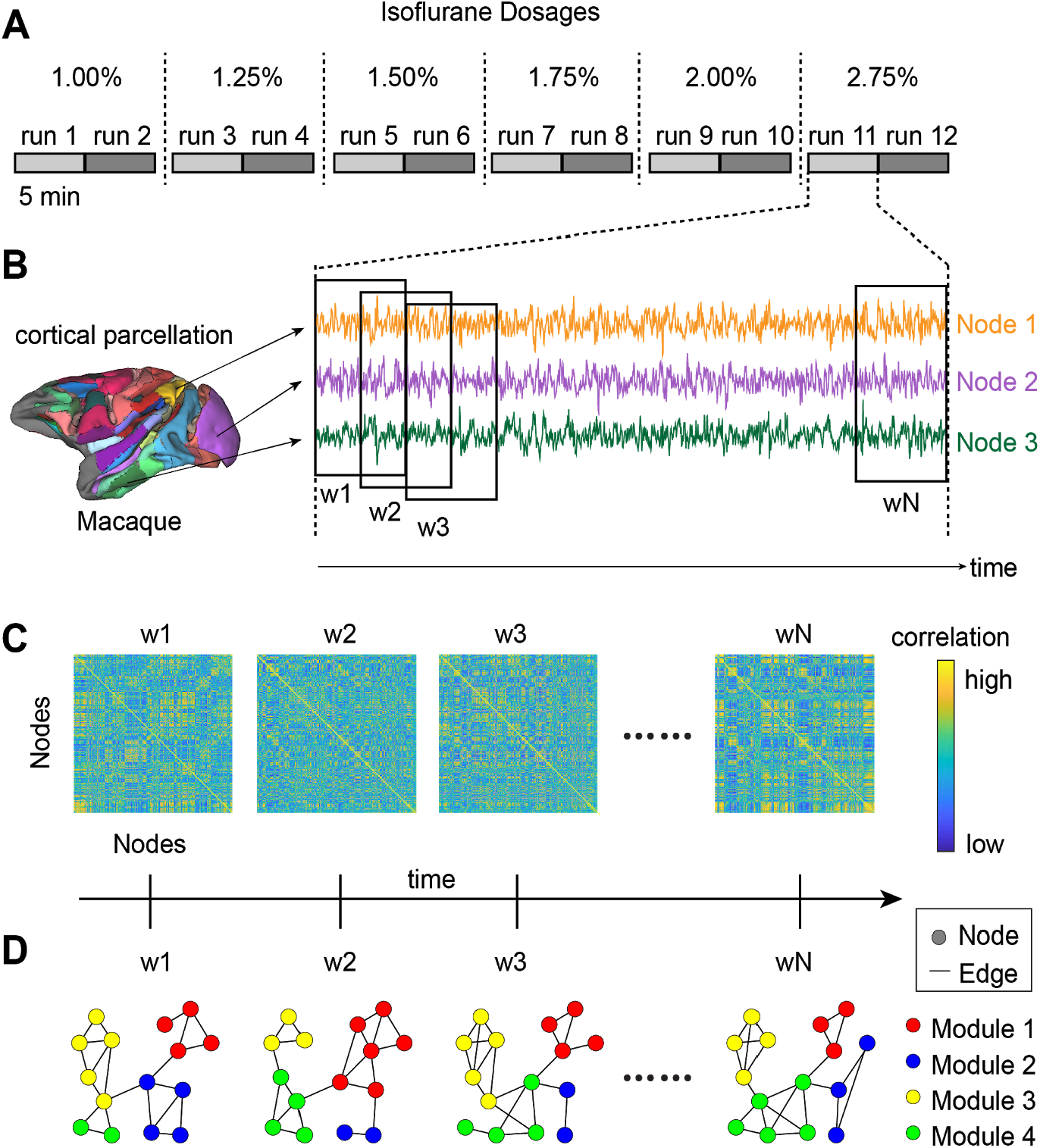
Overview of experiment and analysis approach. **(A)** For each animal, two 5-minute resting-state scans were collected at each of the six isoflurane dose levels. **(B)** Each animal’s cerebrum was parcellated into 174 discrete brain regions and the average BOLD time series was extracted from each region (3 example regions shown). **(C)** The pearson correlation coefficient was calculated for each pair of regions in sliding, half-overlapping windows (shown in B), resulting in whole-brain functional connectivity (FC) matrices for each window (w1 - wN). Together, these FC matrices can be used to construct a multi-slice (temporal) network for each subject and scan (see D). **(D)** Time-resolved clustering was then used to detect temporally dynamic modules in these networks (e.g. four modules in this schematic).

## RESULTS

To examine the effects of general anesthesia on whole-brain temporal modular structure following the induction of unconsciousness, we collected rs-fMRI data from five nonhuman primates at increasing levels of isoflurane dose at 7T MRI. To examine the incremental, dose-dependent effects of isoflurane on temporal module dynamics, we administered isoflurane in six stepwise increments: 1.00%, 1.25%, 1.50%, 1.75%, 2.00% and 2.75% (see Fig. 1A for an overview of our approach). We terminated the session for one monkey that experienced abnormal breathing patterns when the isoflurane concentration was increased to 2.75%, so no data were acquired from that monkey at that particular dose.

Following the parcellation of each animal’s cerebrum into discrete brain regions, the time series correlation was computed between each pair of regions in sliding, half-overlapping windows, resulting in functional connectivity (FC) matrices for each window. We then constructed mutli-slice networks from these FC matrices and partitioned them into temporal modules that maximised a quality function Q (Mucha et al. 2010) (see Fig. 1C). The same measure was calculated for three established classes of null networks (Mucha et al. 2010; Bassett et al. 2011), verifying that subjects’ whole-brain networks exhibited significant modularity (see Supplementary Figure S2).

Statistical analysis of these data is limited by the small number of subjects and the non-independence of the data within each subject (Lazic 2010). Hierarchical modelling is an ideal solution to this problem, but models allowing for subject-specific effects are not easily fit to data sets as small as this one. We therefore used simple linear regression to model network statistics as a function of dose, which can be interpreted as assuming that all subjects respond identically to isoflurane. These models provided good fits overall, as inter-subject variability was generally small relative to dose effects.

### Temporal networks become more fragmented with increasing anesthetic dose

To test the hypothesis that time-resolved modular structure becomes more fragmented with increasing levels of isoflurane dose, we calculated the mean number of modules over all partitions for each subject and scan (two scans per dose). The fit of a linear regression model to these data revealed a positive linear relationship between dose and number (*β*_dose_=0.988, t(56)=5.096, R^2^=0.317, p=4.236e-6; see Figure 2A). We also examined the relationship between dose and the quality function Q, capturing the degree to which whole-brain structure exhibits strong intra-module connectivity and sparse inter-module connectivity, i.e. higher Q corresponds to more isolated modules. A linear regression model revealed a similar relationship with dose as the number of modules (*β*_dose_=0.039, R^2^=0.231, t(56)=4.107, p=1.32e-4; Figure 2B). Together, these findings suggest that deeper levels of sedation partition the brain into a larger number of more isolated functional networks.

**Figure 2:**
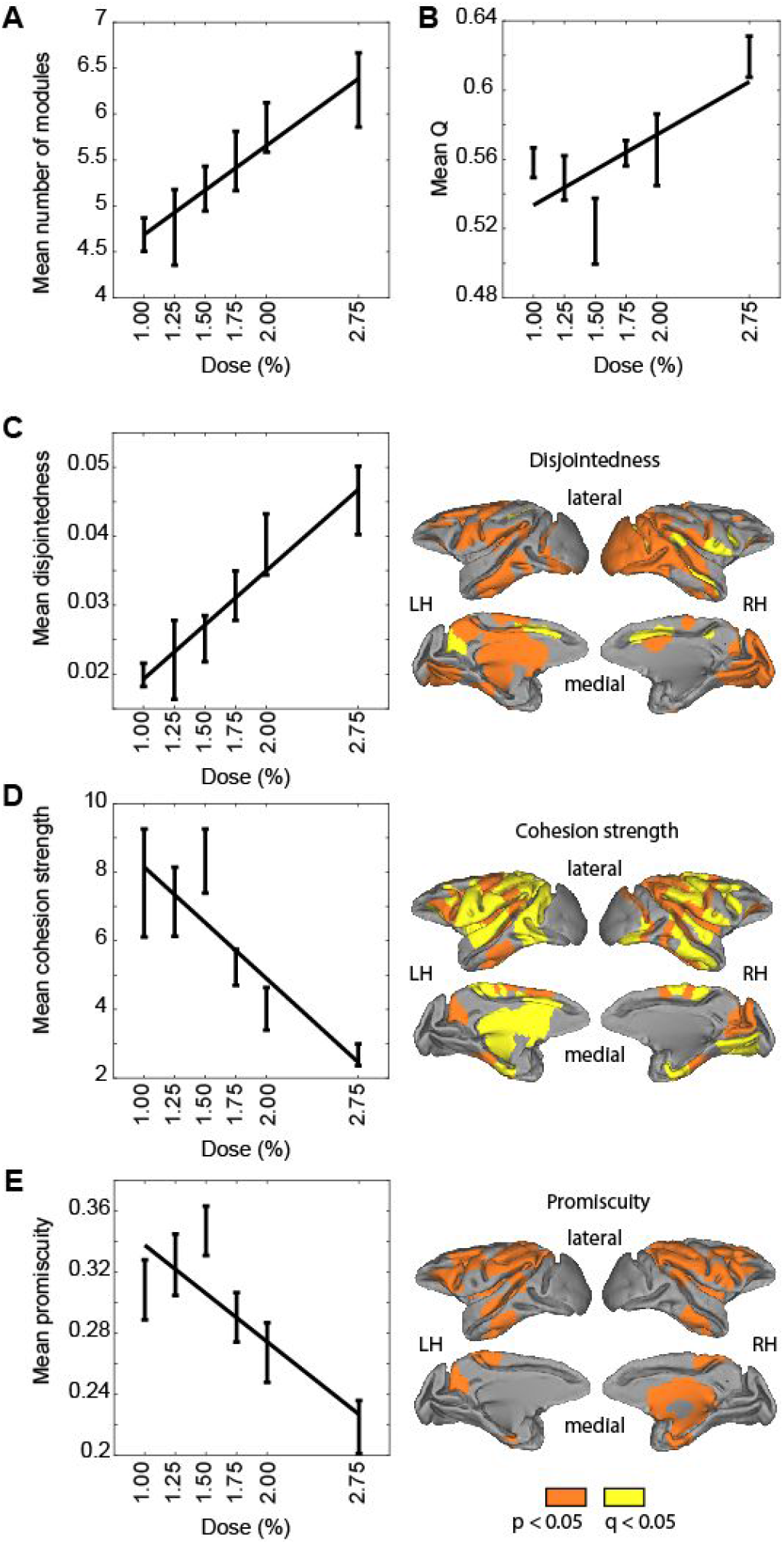
Isoflurane dose affects key properties of whole-brain temporal modules. **(A)** Mean number of temporal modules plotted as a function of isoflurane dose. In the plots, means were first taken over both scans for each dose, and then taken over subjects. Error bars show +/− 1 standard error of the mean. Line shows the fit of a linear regression model to the across-subject means. **(B)** Mean Q, **(C)** mean disjointedness, **(D)** mean cohesion strength and **(E)** mean promiscuity, calculated as in A. Right-side brain plots in panels C, D and E show brain regions for which the relationship (Pearson correlation coefficient) between dose and the statistic on the left had a corresponding p<0.05 (orange regions). Regions in yellow indicate those brain regions at p<0.05 that further passed a false-discovery rate correction (Benjamini and Hochberg 1995).

### Whole-brain reconfiguration becomes more uncoordinated with increasing anesthetic dose

To test our hypothesis that brain regions change modules in a more uncoordinated manner with increasing levels of isoflurane dose, we calculated the disjointed flexibility (disjointedness) of each brain region on each partition, defined by the number of times the region changes modules independently (i.e., without other brain regions) relative to the total number of possible changes (Telesford et al. 2017). We calculated the mean disjointedness by taking the average over all regions on each partition, before taking the mean over these partition means (this procedure was also performed for all other region-specific measures; see below). The fit of a linear regression model revealed a positive linear relationship between dose and disjointedness (*β*_dose_=0.016, t(56)=5.422, R^2^=0.344, p=1.295e-6; Figure 2C). Based on this finding, we hypothesized that the opposite arrangement would also be true, that is, that brain regions would change modules in a more coordinated manner at lower levels of dose. To test this hypothesis, we calculated the strength of cohesive flexibility (cohesion strength) of each region on each partition, an independent measure from disjointedness (Telesford et al. 2017). We calculated this measure by first determining the number of times each region changes modules together with each other region (relative to the total number of possible changes) and then summing over all other regions (Telesford et al. 2017). The fit of a linear regression model revealed a negative linear relationship between dose and cohesion strength (*β*_dose_=-3.276, t(56)=-5.648, R^2^=0.363, p=5.644e-7; Figure 2D). These complementary findings suggest that coordinated modular reconfiguration is a property of whole-brain dynamics under light sedation (and perhaps also a property of consciousness) and that this property breaks down at deeper levels of sedation, such that smaller perturbations (e.g., noise) are sufficient to drive haphazard modular changes.

### Modular reconfiguration becomes more constrained with increasing anesthetic dose

To further characterise modular changes as a function of dose, we calculated the promiscuity of each brain region, defined by the number of modules in which the region participates at least once, relative to the total number of modules (Sizemore and Bassett 2018). The fit of a linear regression model revealed a negative linear relationship between dose and promiscuity (β_dose_=-0.062, t(56)=-5.294, R^2^=0.334, p=2.068e-6; Figure 2E). This finding suggests that despite the fragmentation of whole-brain networks at deeper levels of sedation (Fig. 2A-B), brain regions do not participate in a broader repertoire of these modules.

Taken together, the above findings provide strong evidence that at deeper levels of sedation, (1) whole-brain networks become more fragmented and isolated, (2) individual brain regions move between modules in a more haphazard, uncoordinated manner, and (3) this haphazard movement between a larger number of modules does not entail more diverse modular participation by individual brain regions. Importantly, these results were highly robust across window sizes, the overlap between windows, and the resolution of time-resolved clustering (see Supplementary Figure S3).

### Dynamic network architecture is altered by increasing anesthesia

To investigate whether isoflurane dose influences the degree to which groups of brain regions preferentially interact with one another, we determined the proportion of modular partitions (across all time windows of all scans for all subjects) in which each pair of brain regions was placed in the same module, referred to as a module allegiance matrix (Bassett et al. 2015). This matrix provides a summary of the brain network architecture associated with our isoflurane protocol. We then clustered the module allegiance matrix using symmetric nonnegative matrix factorization (see the Supplementary Methods and Figure S4), identifying four clusters of regions corresponding to whole-brain networks. Because of the known degeneracy of the generalized Louvain algorithm [very different partitions can have nearly identical quality function scores (Bassett et al. 2011)], this clustering approach effectively identifies a consensus modularity partition (Bassett et al. 2013). In other words, it identifies a set of static brain networks summarising the structure of the temporally dynamic modules from all partitions.

This summary network architecture and the module allegiance matrix from which it was derived are shown in Figure 3A-B. For comparison, Figure 3C shows the module allegiance matrix calculated for each level of isoflurane dose (across all time windows of both scans for all subjects, for each dose), labelled according to the four summary networks. Network 2, composed of regions in visual cortex, dorsal parietal and primary somatomotor cortex (yellow in Fig. 3B), appears to remain present across all six levels of dose (Fig. 3C), while the other networks (1, 3 and 4) appear to dissipate with increasing dose. Additionally, whereas the other networks appear to lose their distinctiveness from one another with increasing dose, Network 2 becomes increasingly isolated, and even appears to fracture along hemispheric lines at the highest levels of dose (e.g., 2.00 and 2.75% isoflurane, see Fig. 3D).

**Figure 3:**
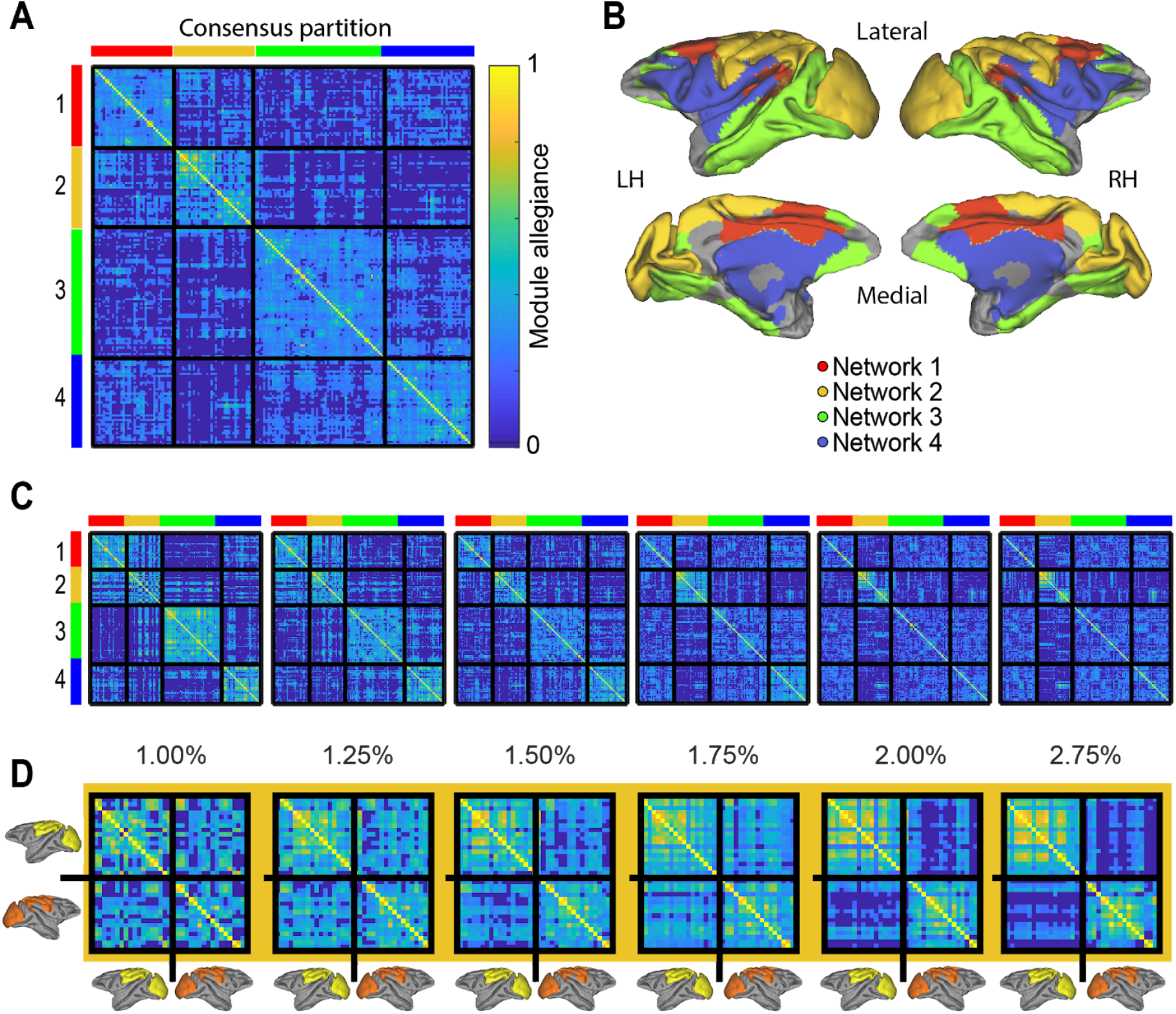
Summary architecture across levels of isoflurane dose. **(A)** Module-allegiance matrix showing the probability of any two brain regions being part of the same temporal module across all subjects, scans and time windows. **(B)** Clustering of the module allegiance matrix in A identified a summary network architecture from the temporal modules, consisting of a cingulate-temporal-parietal-frontal network of regions (Network 1, red), a visual-somatomotor network (Network 2, yellow/orange), a temporal-parietal-prefrontal network (Network 3, green) and a lateral parietal-frontal cingulate temporal network (Network 4, blue). **(C)** The module allegiance matrix for each dose level, labelled according the summary network architecture in A. Note the relatively stable strength of allegiance in Network 2 across doses, compared to the decrease in allegiance in Networks 1, 3 and 4. **(D)** Magnified views of the visual-somatomotor network (Network 2) from C, which highlights a fracturing of this network along hemispheric lines at the highest dose levels (e.g., 2.00 and 2.75% isoflurane). The brain insets with yellow and orange regions denote the labelling of the module allegiance matrix with respect to the left and right hemispheric components of Network 2, respectively.

### Within- and between-network integration is modulated by anesthetic dose

To quantify our observations of the module allegiance matrix, we estimated the integration of the four summary networks, within each network and between networks. With each brain region assigned to a network, the interaction between any two networks can be measured by *I*_*k*1,*k*2_ = (Σ*i* ∈ *C*_*k*1_, *j* ∈ *C*_*k*_1*P_ij_*)/(|*C*_*k*1_||*C*_*k*2_|) (Bassett et al. 2015), where *C*_*k*∈1,2_ are modules, |*C_k_*| is the number of regions they contain, and *P_ij_* is the proportion of the time region *i* and *j* are in the same module. The interaction of a module with itself is calculated by allowing *k*1 = *k*2. The integration between two modules *k*1 ≠ *k*2 is the normalised interaction between them 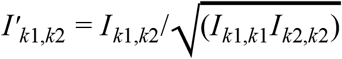. We refer to the interaction of a network with itself as within-network integration and to the integration between different networks as between-network integration. Note that ‘within-network integration’ was referred to as ‘recruitment’ in the task-based analysis from which we borrowed this method (Bassett et al. 2015).

To quantify our observation that the majority of networks (all except the visual-somatomotor network) dissipated at higher levels of isoflurane dose, we fit a linear regression model to within-network integration as a function of dose for all subjects and scans. The model revealed a negative linear relationship between dose and within-network integration (Figure 4A) for Network 1 (*β*_dose_=-2.368, t(56)=-6.3, R^2^=0.415, p=4.92e-8), Network 3 (*β*_dose_=-4.238, t(56)=-6.078, R^2^=0.397, p=1.135e-7) and Network 4 (*β*_dose_=-1.964, t(56)=-4.52, R^2^=0.267, p=3.252e-5). Network 2, the visual-somatomotor network, did not show this relationship (*β*_dose_=-0.217, t(56)=-0.533, R^2^=-0.013, p=0.596). These findings confirm our observation that three of the four networks dissipated at deeper levels of sedation.

**Figure 4:**
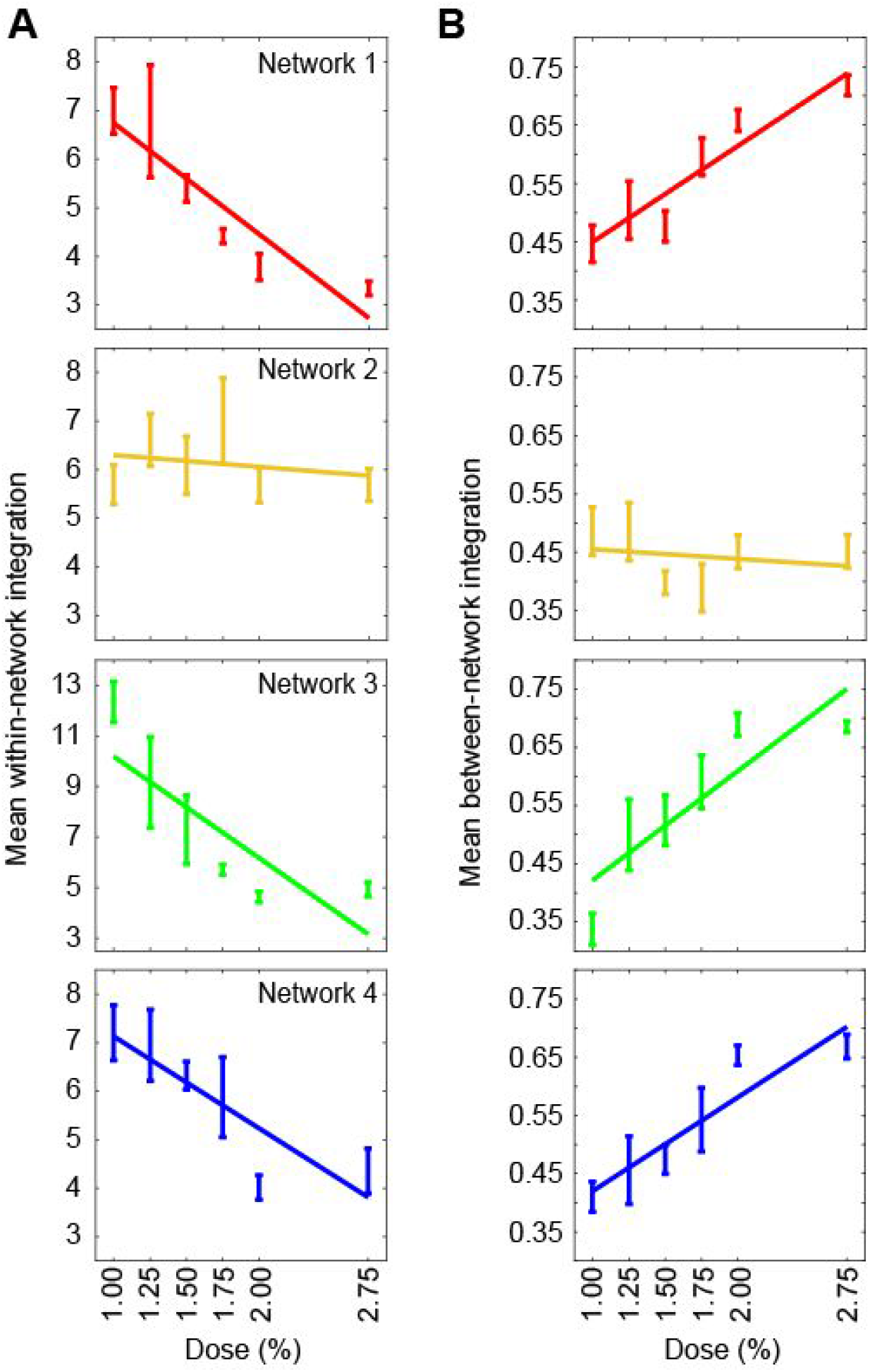
Within- and between-network integration is modified by isoflurane dose for all networks except for the visual-somatomotor network. **(A)** Within-network integration and **(B)** mean between-network integration of each network for each level of isoflurane dose. In the plots, means were taken over both scans for each dose, and then over all subjects. Error bars show +/− 1 standard error of the mean. Lines show fits of a linear regression model to the across-subject means.

To quantify our observation that the majority of brain networks became less distinct with increasing isoflurane dose, we calculated the mean integration between each network and the other three networks. The fit of a linear regression model revealed a positive linear relationship between dose and mean between-network integration (Figure 4B) for Network 1 (*β*_dose_=0.168, t(56)=7.559, R^2^=0.505, p=4.118e-10), Network 3 (*β*_dose_=0.196, t(56)=7.151, R^2^=0.477, p=1.951e-9) and Network 4 (*β*_dose_=0.166, t(56)=6.64, R^2^=0.441, p=1.357e-8). Again, Network 2 was the exception, showing no such dose-dependent relationship with the other networks (*β*_dose_=-0.02, t(56)=-0.743, R^2^=0.01, p=0.46). Furthermore, the integration between each pair of networks precisely tracked these mean interactions, i.e. the integration between Networks 1, 3 and 4 showed a positive linear relationship with dose, whereas the integration between Network 2 (the visual-somatomotor network) and each of the other networks failed to show a dependence on dose (see Supplementary Figure S5). Together, these findings confirm our observation that three out of four networks became less distinct with increasing isoflurane dose.

### Network-specific effects on region-based measures of modular reconfiguration

Lastly, having found network-specific differences for within-network and between-network integration, we sought to determine whether the movement of brain regions between temporal modules would also show differences according to network membership. We therefore calculated the mean disjointedness, cohesion strength and promiscuity of each summary network, taking the same approach as above (Figure 2C-E), but averaging these measures within each network, rather than across all brain regions. For each network, we fit a linear regression model to each measure of modular adaptability as a function of dose, finding that mean disjointedness increased with dose for all networks (Figure 5A, left), and that mean cohesion strength (Figure 5B, left) and promiscuity (Figure 5C, left) decreased with dose for all networks. Across the twelve fits, R^2^ values ranged from 0.173 to 0.384 and p-values ranged from 0.01e-3 to 2.13e-7. Averaging across dose levels for each subject, the respective magnitudes of disjointedness (Figure 5A, right) and promiscuity (Figure 5C, right) were notably smaller in the visual-somatomotor network, while the magnitude of cohesion strength did not obviously differ between networks (Figure 5B, right). Together, these findings suggest that the increase in uncoordinated modular reconfiguration at higher levels of sedation occurs globally throughout the brain, but that the magnitude of the effect is smaller in primary sensory and motor areas (which make up the visual-somatomotor network).

**Figure 5:**
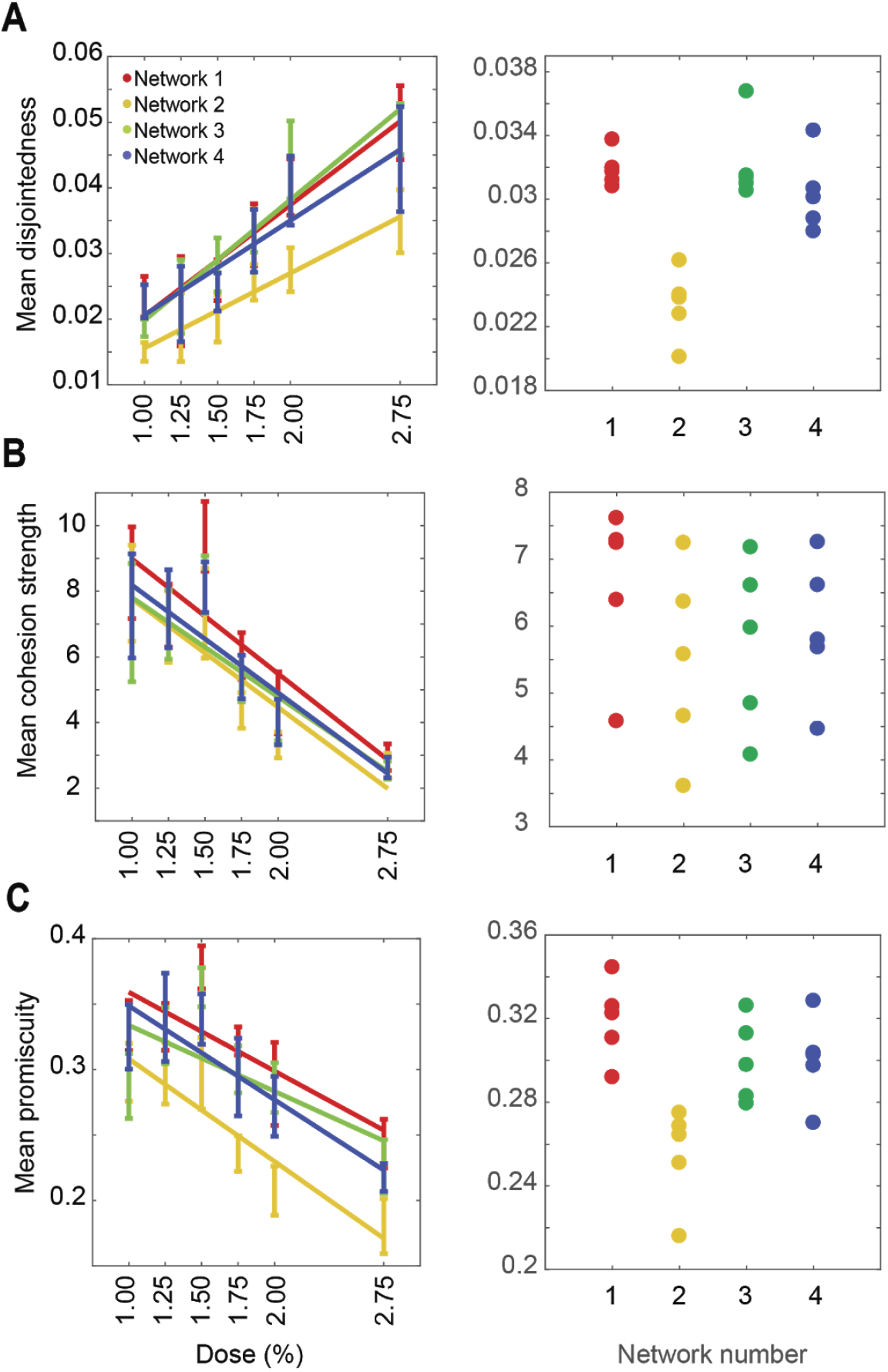
Mean disjointedness, cohesion strength and promiscuity of each network. **(A)** Mean disjointedness for each level of isoflurane dose (left), where means were taken over both scans for each dose, and then over all subjects. Error bars show +/− 1 standard error of the mean. Lines shows fits from a linear regression model to the across-subject means. Distribution of subject means, taken over all doses is shown on the right for each network. (B) Mean cohesion strength. (C) Mean promiscuity.

## DISCUSSION

We investigated the dose-dependent effects of isoflurane on temporal modular structure in nonhuman primates following the induction of unconsciousness. Our analyses revealed that whole-brain structure became more fragmented at deeper levels of sedation, where the number and isolation of temporal modules increased with dose (Figure 2A-B). When we characterized this modular reconfiguration at the level of brain regions, we found that deeper levels of sedation were associated with more uncoordinated movement of brain regions between modules, as revealed by an increase in disjointed flexibility (Figure 2C) and a decrease in cohesive flexibility (Figure 2D). Notably, this uncoordinated reconfiguration coincided with a proportional decrease in the number of modules in which brain regions participated, as measured by their promiscuity (Figure 2E). Next, by determining the probability that each pair of brain regions was assigned to the same module over time, we identified four whole-brain networks that summarised subjects’ dynamic whole-brain architecture across levels of sedation (Figure 3A-B). Three out of four of these networks dissipated at deeper levels of sedation, as measured by their within-network (Figure 4A) and between-network (Figure 4B) integration. Interestingly, a lone network comprised of visual and somatomotor regions was impervious to these dose-dependent effects on integration. Together, our findings indicate that higher anesthetic dose results in the uncoordinated reconfiguration of modular structure across cortex, but that the breakdown in network structure is relatively spared in primary visual, somatosensory and motor regions. Consistent with recent theoretical work (Tononi et al. 2016), these dose-dependent effects on whole-brain modular structure suggest that unconsciousness may be graded in nature, and driven by disordered communication between circuits within association cortex. These findings (summarised in Figure 6) not only characterise the dynamics of whole-brain network structure across depths of unconsciousness, but they further illuminate the effects of anaesthetic dose in clinical treatment, and may offer diagnostic tools for identifying residual consciousness in vegetative state patients (Owen et al. 2006; Sitt et al. 2014).

**Figure 6:**
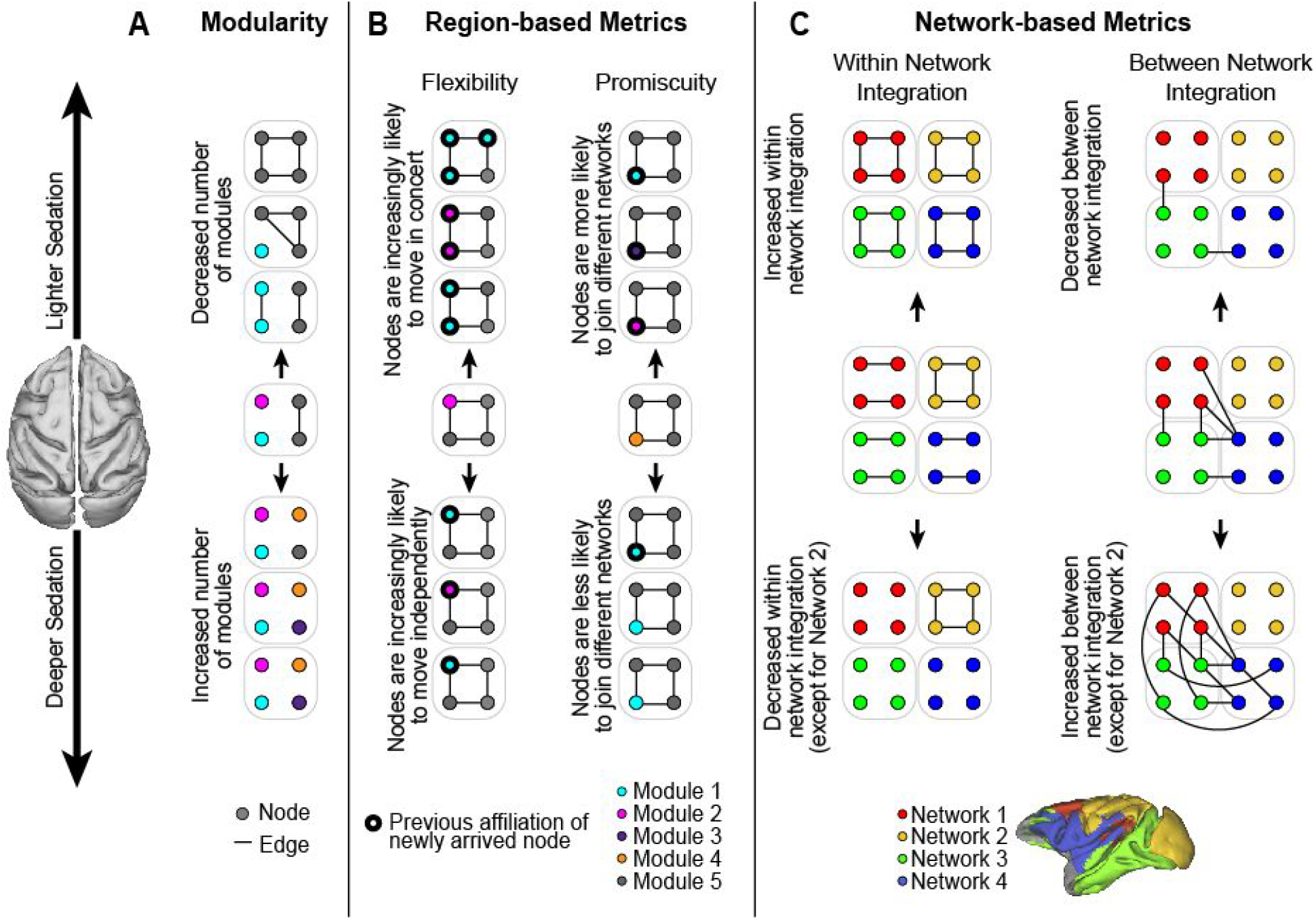
Summary of results. **(A)** A larger number of more isolated modules were detected at higher levels of anaesthetic dose, quantifying an increase in whole-brain fragmentation at deeper levels of unconsciousness. **(B)** Modular flexibility was less coordinated (more disjointed) at higher levels of dose, with brain regions participating in a smaller proportion of modules on average (lower promiscuity). **(C)** With the exception of a visual-somatomotor network (yellow), summary networks derived from the temporal modules became less distinct at higher levels of dose, quantified by their within-network and between-network integration.

### Current findings in the context of past work

Several earlier studies have used static module detection methods to characterise network fragmentation during unconsciousness (Achard et al. 2012); (Boly et al. 2012; Spoormaker et al. 2012; Monti et al. 2013; Tagliazucchi et al. 2013; Hutchison et al. 2014), and so it is important to differentiate the present methodology from that of these earlier investigations. Static module detection methods operate according to the same general principle as temporal module detection methods (i.e. they partition a network into modules that typically maximize the ratio of within-module to between module connectivity), but they do so for single-layer networks. Thus, even if these methods are used in each layer of a multi-layer network constructed from windowed time series [e.g. (Tagliazucchi et al. 2013)], there is no formal relationship between a module in one time window and the modules in any other time window [see (Mucha et al. 2010; Bassett et al. 2011)]. Consequently, summary statistics (e.g., Q and the number of modules) can be computed over multiple time windows, but modular reconfiguration cannot be measured. Our temporal module analysis was in large part motivated by these earlier studies, which provided evidence for the increased fragmentation of whole-brain networks according to the number of static modules during isoflurane- (Monti et al. 2013; Hutchison et al. 2014) and propofol-induced (Monti et al. 2013) unconsciousness, as well as the magnitude of static modularity during propofol-induced unconsciousness (Monti et al. 2013) and sleep (Boly et al. 2012; Spoormaker et al. 2012; Tagliazucchi et al. 2013). Our finding that both Q (the magnitude of modularity, Figure 2B) and the number of temporal modules (Figure 2A) increased with dose offers novel support for the hypothesis that network fragmentation occurs in a graded fashion across levels of sedation. In this regard, it is important to note how these two measures are distinct. Q captures the degree to which modules are isolated from one another, but does not imply that more (or less) modules exist in a given partition. An increase in the number of modules with dose indicates a different kind of fragmentation altogether. Concurrently, these two findings further characterise a large body of evidence from static FC analyses that shows the breakdown of distributed networks, whereby brain networks decompose into a larger number of more isolated sub-networks during unconsciousness [see (MacDonald et al. 2015; Cavanna et al. 2018)].

It is also important to note that maximal modularity does not equate to optimal modularity. While modules confer functional specialisation and robustness, excessive modularity foregos the advantages of integration over specialised subsystems (Kirschner and Gerhart 1998; Kashtan and Alon 2005; Wagner 2013). Thus, optimal modularity can be defined as a balance between these competing requirements. Our results imply that this balance is increasingly skewed toward specialisation at deeper levels of unconsciousness, and we envision a continuum of optimal balance that peaks at higher levels of alertness during conscious processing. From this viewpoint, it is intuitive that connectivity between association cortical areas breaks down at lower levels of sedation than connectivity between sensory and motor systems (see below). Integration over specialised subsystems is widely believed to be the role of association cortex in the cortical hierarchy (Jones and Powell 1970; Kaas 1989), supporting the analytic and creative processing associated with awareness and higher cognitive function.

Optimal modularity bears an intriguing similarity to the foundational principles of integrated information theory, which posit that consciousness emerges from a balance between differentiation and integration of distributed computational components (Tononi et al. 2016). This balance is hypothesised to maximise systemic complexity, resulting in a large number of more diverse brain states, required to account for a large, diverse set of conscious experiences (Cavanna et al. 2018). Our finding that fragmentation (corresponding to differentiation) increased with dose appears to support this hypothesis. Future work should systematically explore the relationship between measurements of modularity and complexity in whole-brain networks, and the dependence of this relationship on depths of unconsciousness.

The dose-dependent decrease in within-network integration in three out of four summary networks derived from temporal modules (Figure 4A) accords with earlier evidence for the breakdown of static networks at deeper levels of sedation [see (MacDonald et al. 2015)], while the imperviousness of the visual-somatomotor network to this effect provides further evidence that the breakdown of network structure under supra-threshold anaesthetic dose is spatially non-uniform, with primary sensory and motor cortical areas less affected than higher-order association cortical areas (He et al. 2008; Liu, Zhu, et al. 2013; Hudetz et al. 2016). At first glance, our finding that between-network integration across the majority of networks increased with dose (Figure 4B) appears to conflict with earlier evidence showing a *decrease* in the integration between networks under anaesthesia (see MacDonald et al. 2015). However, we measured network integration relative to a module allegiance matrix, which describes the probability that regional pairs were assigned to the same module over time (Figure 3A), not the magnitude of coactivation of their constituent regions (as is typical in static FC investigations). To reconcile these observations, we calculated the mean FC of our summary networks and found that it decreased with dose, both within and between networks (see Supplementary Figure S6A,C). Thus, the increase in between-network integration among three of four summary networks reflects a general weakening of network structure at higher levels of dose, rather than an increase in functional interactions. In addition, the preservation of the integrity of the visual-somatomotor network across levels of dose may reflect the overall reduction in FC, which tends toward the underlying structural connectivity (see MacDonald et al. 2015).

Our finding that disjointed flexibility increased with isoflurane dose (Figure 2B) supports our hypothesis that weaker network structure should lead to more haphazard, uncoordinated changes in the affiliation of brain regions with modules. Likewise, our finding that cohesion strength decreased with increased isoflurane dose (Figure 2C) is consistent with the decrease in whole-brain state transitions at higher dose (Hutchison et al. 2014; Barttfeld et al. 2015), assuming that such state transitions are more readily driven by groups of brain regions acting in concert than by individual brain regions. The inverse relationship between disjointedness and cohesion strength, which serve as two independent measures of regional flexibility (Telesford et al. 2017), suggests that these measures may also provide a useful marker for tracking levels of conscious processing. Prior work using dynamic connectivity methods has shown that the awake state is characterized by a rich and flexible repertoire of whole-brain states (Barttfeld et al. 2015; Uhrig et al. 2018), and that these states are expressed more frequently at lighter (compared to deeper) levels of sedation (Hutchison et al. 2014; Barttfeld et al. 2015). From this perspective, coordinated changes in modular structure may provide a signature of an engaged mind, suggesting that cohesive flexibility should correlate with the performance of demanding cognitive tasks, from learning (Telesford et al. 2017) and problem solving, to creative thinking. Our finding that promiscuity increased with decreasing isoflurane dose is also consistent with this possibility, assuming that the exploration of a larger number of diverse brain states entails a broader range of modular reconfigurations, and correspondingly, a proportionally larger repertoire of modules in which brain regions participate. Future work should address these possibilities.

### Methodological Considerations

Our findings should be interpreted in light of several methodological considerations. Firstly, our study did not measure subjects’ FC during the awake state, restricting our discussion to changes in network architecture during unconsciousness and at deepening levels of sedation. Prior work with these data showed a stepwise, gradual decrease in the repertoire of dynamic FC states with increasing dose, while the stability of each of these states increased, as measured by the number of state transitions (Hutchison et al. 2014). Another recent study with nonhuman primates used propofol anaesthesia to characterize the transition between consciousness and unconsciousness, replicating this general pattern of results and showing that the predominant FC brain state during unconsciousness closely resembled the underlying anatomical map. This finding has been further demonstrated in both rats under isoflurane anesthesia (Ma et al. 2017) and in human deep sleep (Tagliazucchi et al. 2016), suggesting that anesthesia serves to constrain the dynamic repertoire of FC patterns onto an anatomical structural backbone (see also Fischer et al. 2018). As alluded to above, our findings provide a complementary perspective from which to view these results: the type of modular flexibility expressed by brain regions at different levels of sedation (e.g., disjointedness vs. cohesive flexibility) may ultimately underly the phenomenological effects of anesthesia.

Secondly, unconsciousness in our study was induced through isoflurane, a commonly used anesthetic in rs-fMRI investigations in nonhuman primates and rodents (Vincent et al. 2007; Shmuel and Leopold 2008; Hutchison et al. 2010; Mars et al. 2011; Liu, Zhu, et al. 2013). As such, it is possible that our results are primarily attributable to its specific mechanisms of action. Isoflurane is a preferred anesthetic in animal studies due to its robustness, relative ease of use, low cost, high survivability, and speed of induction and recovery (see also Hutchison et al. 2014). It operates through the targeting of multiple membrane channels and receptor types (reviewed in Franks 2008), and its administration is characterized by slow-wave activity in electroencephalographic (EEG) recordings (Nickalls and Mapleson 2003). At levels above 1 MAC [the concentration of anesthetic needed to prevent motor responses in 50% of subjects in response to surgical stimulus (typically an incision)] burst suppression occurs, characterised by periods of high amplitude neural activity (bursts) that alternate with isoelectric quiescence (suppression). These periods of suppression are enhanced at increased levels of dose, culminating in electrical silence at 2-2.5 MAC (Hoffman and Edelman 1995). The isoflurane levels used here (1.00-2.75%) correspond to a range of 0.78-2.15 MAC, suggesting that our isoflurane protocol permitted sampling of the entire transition from continuous low-frequency activity to near complete burst suppression and neural silence.

Thirdly, it is recognized that isoflurane, in addition to having effects on metabolic rate, has effects on cerebral blood flow and blood volume (Masamoto and Kanno 2012). Consequently, there are concerns that neurovascular effects at higher doses in anesthesia-related rs-fMRI investigations obscure potential neural changes. However, studies combining fMRI with electrical recordings, such as EEG-fMRI (Vincent et al. 2007; Liu, Pillay, et al. 2013; Barttfeld et al. 2015; Ranft et al. 2016), have generally indicated a close relationship between changes in electrical activity and changes in hemodynamics and FC. It has also been demonstrated that different types of anesthesia, from propofol and isoflurane to sevoflurane and ketamine, exhibit near-identical effects on the repertoire of functional brain states (Hutchison et al. 2014; Barttfeld et al. 2015; Uhrig et al. 2018). As such, our findings on temporal modularity are unlikely to reflect only the mechanisms of isoflurane. Rather, they are likely to reflect a general cortical signature of anesthesia-induced loss of consciousness.

### Links to current theories on consciousness

Unconsciousness has been extensively characterised as a decrease in the integration of regional activity in whole-brain networks, both theoretically (Dehaene et al. 2014; see Tononi et al. 2016) and experimentally (MacDonald et al. 2015; see Cavanna et al. 2018)]. Two general mechanisms have been suggested to underlie this decrease (Hudetz 2006; Alkire et al. 2008). On the one hand, FC breaks down non-uniformly at supra-threshold levels of anaesthesia, suggesting an overall decrease in correlated activity between brain regions (Peltier et al. 2005; Lu et al. 2007; Deshpande et al. 2010). On the other hand, a global decrease in functional segregation suggests a decrease in the specificity of connectivity (Liu et al. 2011; Kalthoff et al. 2013; Liu, Zhu, et al. 2013). Our results take an important step toward reconciling these observations, offering evidence in support of both views within a common framework. With respect to the first view, our finding that the integrity of three out of four summary networks diminished at deeper levels of sedation provides compelling evidence for the spatially non-uniform breakdown of network structure. With respect to the second view, our finding that both the strength of modularity and the number of modules increased at deeper levels of sedation implies an increase in functional segregation, rather than a decrease; however, our finding that the strength of FC decreased both within and between networks at deeper levels of sedation (Supplementary Figure S6A,C) is consistent with a general breakdown of network connectivity. Moreover, the dose-dependent increase in disjointedness, decrease in cohesion strength, and decrease in promiscuity occured in all networks (Figure 5A), further demonstrating global network-level effects. Indeed, we hypothesised that the global weakening of FC would render small background perturbations sufficient to drive uncoordinated modular changes (Figure 2B). This weakening of FC and its effects on modular reconfiguration may underlie the dissipation of three out of four summary networks, since our module allegiance matrix quantified the probability that brain regions were grouped together in dynamic modules. Thus, in so far as our temporal modular approach addresses each of the core aspects of the two views, our findings suggest that they account for quantifiable expressions of the same underlying mechanism.

## Acknowledgements

This work was supported by operating grants from the Canadian Institutes of Health Research (CIHR) awarded to J.P.G (MOP126158), S.E. (MOP89785) and R.M. (PRG-165679). D.S. was funded by the European Union’s Horizon 2020 Research and Innovation Programme under the Marie Sklodowska-Curie grant agreement No 798255. C.N.A. was supported by a Natural Sciences and Engineering Research Council (NSERC) graduate award. J.P.G. was also supported by a NSERC Discovery Grant, as well as funding from the Canadian Foundation for Innovation. The authors thank S. Hughes for technical assistance.

