## Supplementary material for "Dynamic reconfiguration, fragmentation and integration of whole-brain modular structure across depths of unconsciousness"

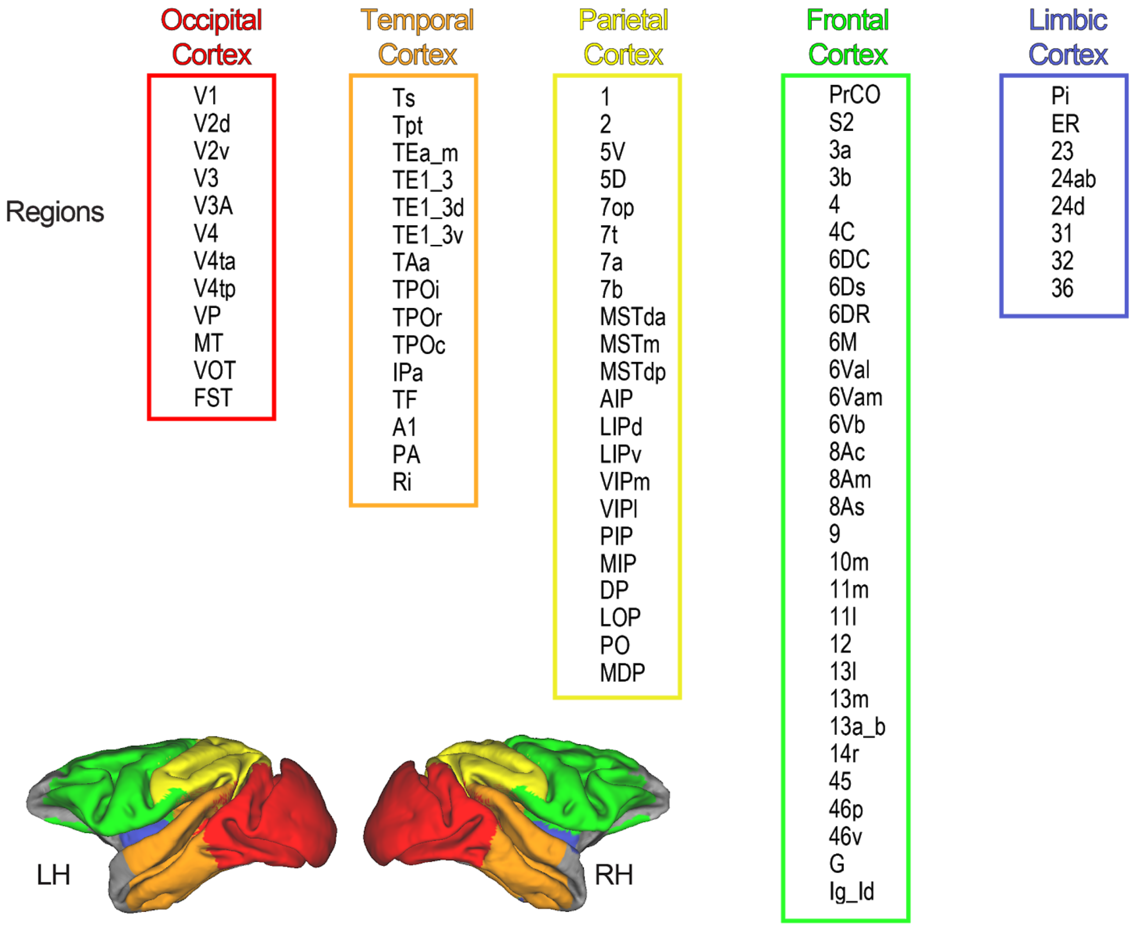


**Figure S1. Cortical regions used for analysis.** Top, Region labels used for temporal module detection analysis, with associated bounding boxes color-coded according to their neuroanatomical location in cortex (Lewis and Van Essen 2000a, 2000b). Bottom, Neuroanatomical locations are displayed visually on the macaque F99-template. LH = left hemisphere; RH = right hemisphere.

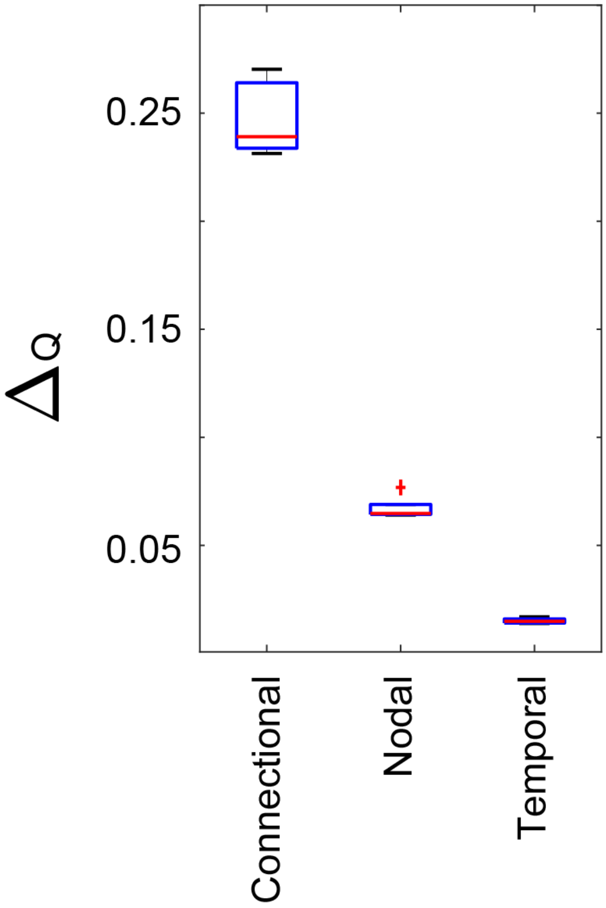


**Figure S2. Subjects' whole-brain temporal networks exhibit significant modular structure.** Boxplots show differences (real minus null) in the modularity quality function (Q) between subjects' networks and permuted networks for connectional (one-sample t-test, d.f.=4, t=31.551, p=6.015e-6), nodal (d.f.=4, t=27.967, p=9.724e-6) and temporal (d.f.=4, t=27.282, p=1.074e-5) null models [see (Bassett et al. 2011)]. Differences were calculated over all 12 scans. Red horizontal lines show medians. Black lines show minimum and maximum values. Bottom and top of blue boxes correspond to 25th and 75th percentiles. Red + symbol indicates outlier.

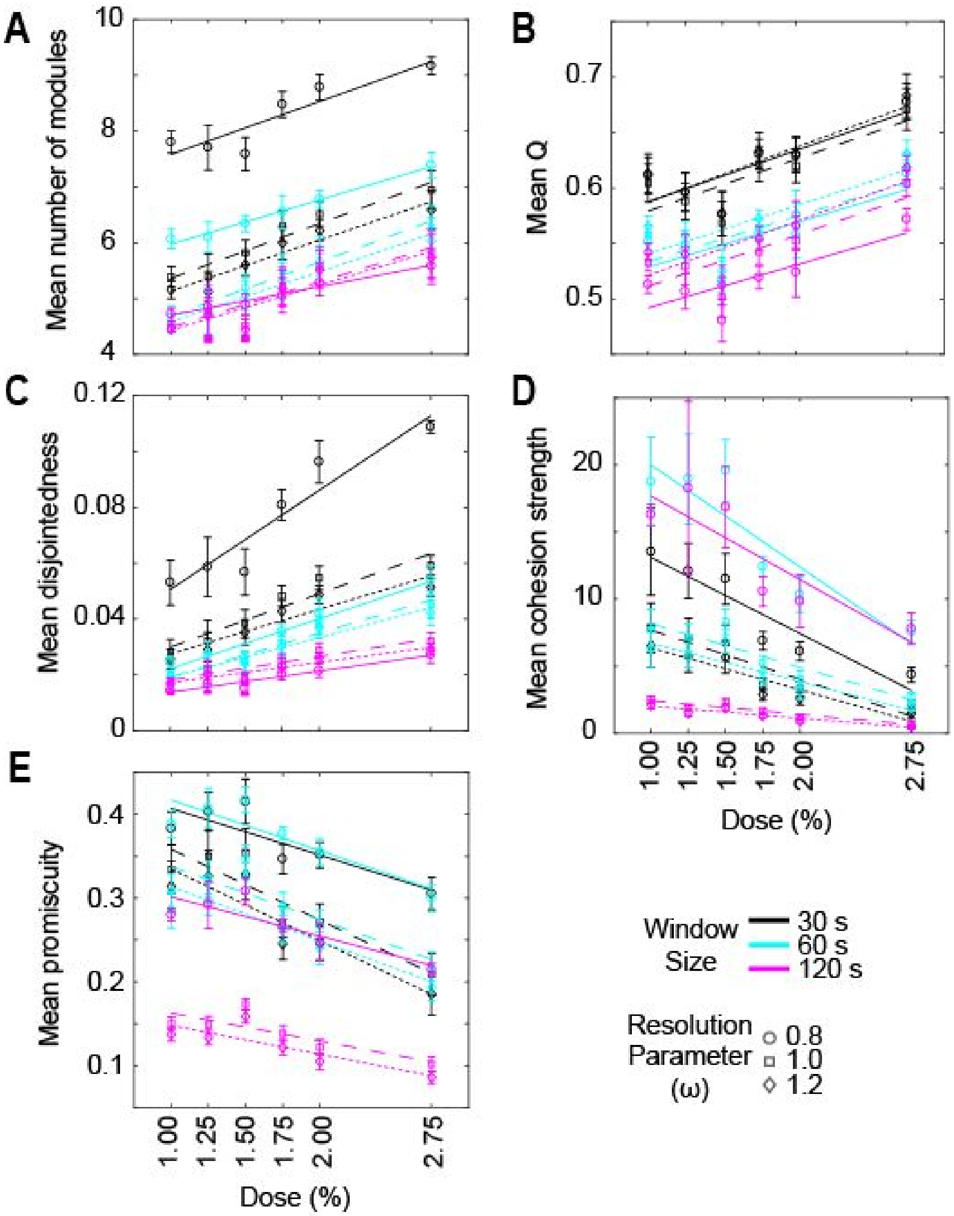


**Figure S3. Our results were robust to changes in the size of the time window used to construct functional networks, the overlap between time windows, and the temporal resolution of time-resolved clustering.** The strong, linear effect of dose on the mean number of modules **(A)**, Q **(B)**, disjointedness **(C)**, cohesion strength **(D)** and promiscuity **(E)** was preserved over three widely varying window sizes (30s, black; 60s, cyan; 120s, magenta) and three values of the resolution parameter $\omega$ of the generalised Louvain algorithm used here (0.8, circles; 1.0, squares; 1.2, diamonds). The step between windows was 30s in all cases, so the overlap between windows was 0%, 50% and 75% for window sizes of 30s, 60s and 120s, respectively. Over the five module statistics and nine combinations of parameter values, the fit of a linear regression model yielded mean R^2=0.304 ($0.162 \le$ R^2 $\le 0.456$) with corresponding mean p=0.1.302e-4 (5.972e-9 $\le p \le$ 0.002). Solid, dashed and dotted fitted lines correspond to $\omega$ values of 0.8, 1.0 and 1.2 respectively. In the plots, means were taken over both scans for each dose, and then over all subjects. Lines show fits of a linear regression model to the across-subject means.

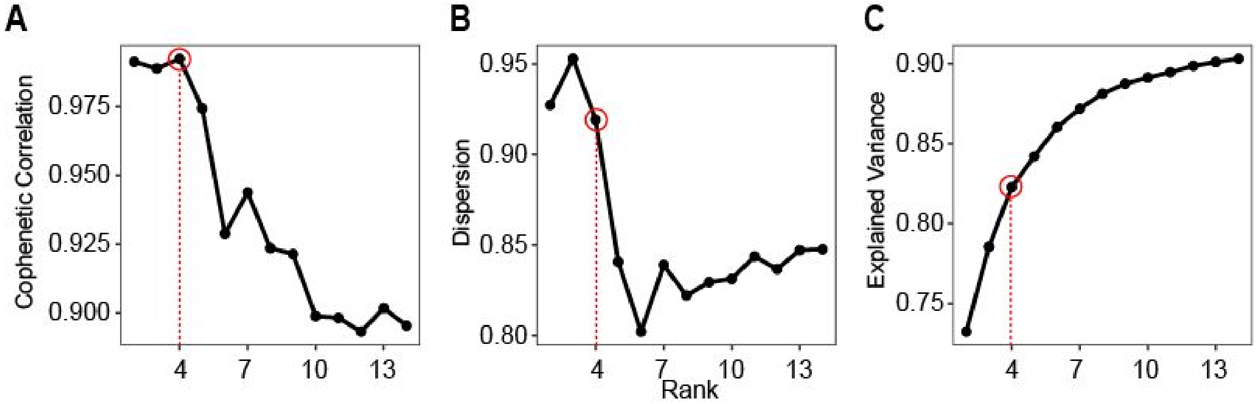


**Figure S4: Clustering diagnostics for symmetric non-negative matrix factorization (SymNMF) of the module allegiance matrix.** The number of clusters was chosen by fitting 250 random initializations of each factorization rank in the range of 2 to 15. For each rank, we computed the average root mean square reconstruction error, the cophenetic correlation coefficient and the dispersion index. We judged that a rank 4 solution was acceptable, as this was sufficient to explain a large majority of the variance in the observed data (over 80%), and both the cophenetic correlation and dispersion index began to decrease sharply beyond this point. We then selected the rank 4 solution with the lowest reconstruction error, and generated a clustering by assigning each brain region to the factor on which it loaded most strongly.

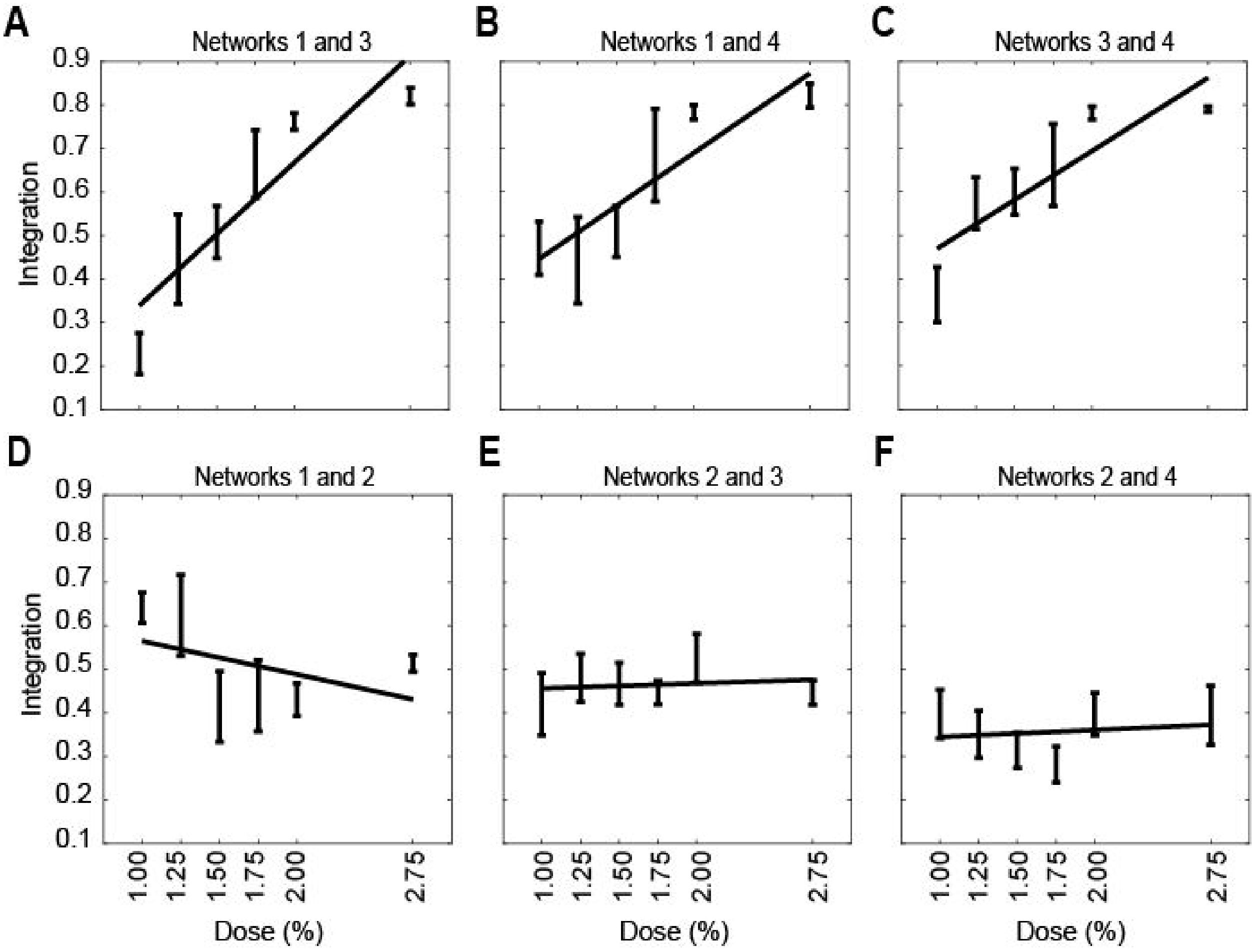


**Figure S5. Between-network integration for each pair of summary networks for each dose level. (A)** The fit of a linear regression model to the integration between Networks 1 and 3 for all subjects and scans showed a positive linear relationship between dose and integration ($\beta_{dose}$=0.34, t(56)=8.388, $R^2$=0.557, p=1.788e-11). **(B)** Integration between Networks 1 and 4 ($\beta_{dose}$=0.25, t(56)=5.652, $R^2$=0.363, p=5.561e-07). **(C)** Integration between Networks 3 and 4 ($\beta_{dose}$=0.233, t(56)=6.3728, $R^2$=0.42, p=3.738e-8). **(D)** Integration between Networks 1 and 2 ($\beta_{dose}$=-0.087, t(56)=-1.9485, $R^2$=0.064, p=0.056) ($R^2$=0.064, p=0.056). **(E)** Integration between Networks 2 and 3 ($\beta_{dose}$=0.015, t(56)=0.47264, $R^2$=0.004, p=0.638). **(F)** Integration between Networks 2 and 4 ($\beta_{dose}$=0.013, t(56)=0.39 , $R^2$=0.003, p=0.698). In the plots, means were taken over both scans for each dose, and then over all subjects. Lines show fits of a linear regression model to the across-subject means.

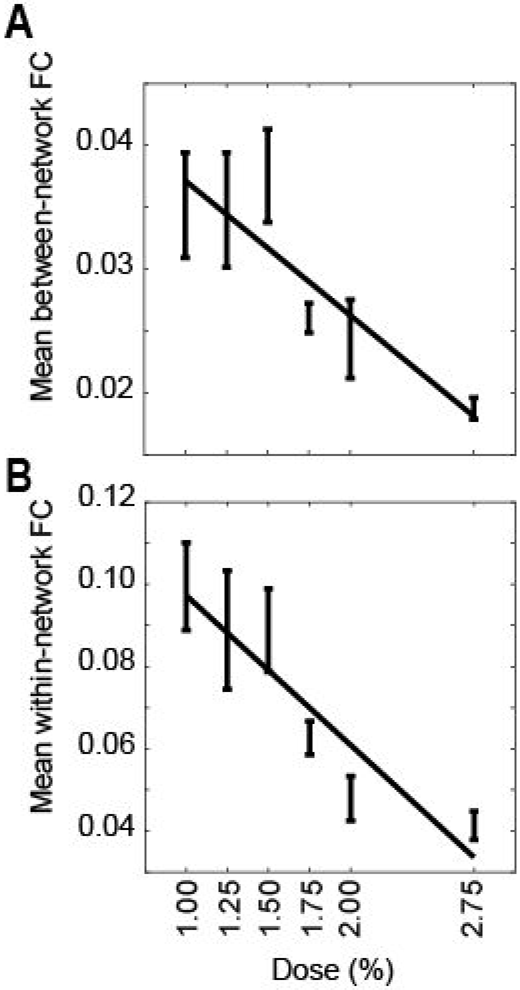


**Figure S6. Between-network and within-network functional connectivity (FC) for each dose level. (A)** The fit of a linear regression model to mean between-network FC for all subjects and scans showed a negative linear relationship between dose and FC, where the mean for each network was taken over its FC with the other three networks, before taking the mean over all networks ($\beta_{dose}$=-0.011, t(56)=-5.212, $R^2$=0.327, p=2.787e-6). **(B)** Mean within network FC showed a negative linear relationship with dose, where means were taken over all networks ($\beta_{dose}$=-0.037, t(56)=-6.745, $R^2$=0.448, p=9.118e-9). In the plots, means were taken over both scans for each dose, and then over all subjects. Lines show fits of a linear regression model to the across-subject means.

### *SUPPLEMENTARY METHODS*

***Clustering of the module allegiance matrix with non-negative matrix factorisation.*** We clustered the module allegiance matrix using symmetric nonnegative matrix factorization [SymNMF; (Kuang et al. 2012)] , which decomposes a symmetric matrix $Y$ as $Y = AA^T$ , where $A$ is a rectangular matrix containing a set of nonnegative factors which minimize a sum-of-squares error criterion. SymNMF has previously shown good performance for factoring similarity matrices (Kuang et al. 2012, 2015), and has the benefit of producing a natural clustering solution by assigning each observation to the factor on which it loads most strongly.

To choose the number of factors, we fit 250 random initializations of each factorization rank in the range of 2 to 15. For each rank, we computed the average root mean square reconstruction error, dispersion coefficient (Kim and Park 2007), and cophenetic correlation (Brunet et al. 2004). The latter two measures quantify the consistency of the clustering solutions returned by different initializations.

We judged that a rank 4 solution was acceptable, as this was sufficient to explain a large majority of the variance in the observed data (over 80%), and both the dispersion and cophenetic correlation began to decrease sharply beyond this point (Figure S4). We then selected the rank 4 solution with the lowest reconstruction error, and generated a clustering solution by assigning each brain region to the factor with the highest loading.
